# Bacterial suppressor-of-copper-sensitivity (Scs) proteins exhibit diverse thiol-disulfide oxidoreductase cellular functions

**DOI:** 10.1101/2023.02.07.527441

**Authors:** Yaoqin Hong, Jilong Qin, Lachlan Mitchell, Jason J. Paxman, Begoña Heras, Makrina Totsika

**Author notes:** Correspondence to Makrina Totsika.

## Abstract

Disulfide bond (Dsb) proteins catalyse oxidative protein folding governing bacterial survival and virulence. Dsb systems in *Escherichia coli* K-12 are well-studied, yet what determines dithiol oxidase or disulfide reductase activity remains unknown. Past studies suggest oligomerisation of periplasmic thiol oxidoreductases dictates the direction of thiol catalytic activity. Here, we studied three suppressor-of-copper-sensitivity C (ScsC) Dsb-like proteins known to exist in the reduced state and bind to copper. These proteins adopt different quaternary structures: *Salmonella enterica* ScsC (StScsC) is monomeric, while ScsC from *Proteus mirabilis* (PmScsC) and *Caulobacter crescentus* (CcScsC) are trimeric. When expressed in the model organism *E. coli* K-12, we showed that all three ScsC proteins exhibit both dithiol oxidation and disulfide reduction activity, despite structural differences. Interestingly, while ScsC reductase function was supported by the canonical *E. coli* DsbD reductase, oxidase activity depended on environmental oxidation. However, an engineered monomeric PmScsC synergises with *E. coli* DsbB to gain dithiol oxidase activity at the expense of reductase function. Thus, oligomerisation could be one mechanism by which ScsC proteins avoid interactions with the periplasmic thiol oxidase pathway. This tightly controls their re-oxidation and maintains ScsC proteins in the reduced state required for binding and sequestering toxic levels of cellular copper.

## INTRODUCTION

Intramolecular disulfide bonds (Dsb) between cysteine (Cys) residues play a central role in the function and stability of many secreted and membrane proteins in both eukaryotes and prokaryotes (Sevier & Kaiser, 2002). In Gram-negative bacteria, oxidative protein folding takes place in the periplasmic space and is catalyzed by a group of redox-active disulfide bond (Dsb) proteins characterised by the presence of a thioredoxin (TRX) domain and a conserved CXXC active site (Heras *et al.*, 2009, Landeta *et al.*, 2018). The *Escherichia coli* K-12 oxidative protein folding machinery includes four Dsb proteins organised in two distinct redox pathways: the DsbA/DsbB oxidative pathway that introduces disulfide bonds between cysteine residues in substrate proteins and the DsbC/DsbD isomerase pathway that rearranges incorrect disulfide bonds introduced in substrate proteins (Manta *et al.*, 2019, Heras *et al.*, 2007, Bardwell *et al.*, 1993). In this system, the oxidase DsbA enzyme is monomeric (Martin *et al.*,1993), whereas the isomerases DsbC (or DsbG) (McCarthy *et al.*, 2000, Heras *et al.*, 2004) are dimeric with their catalytic motifs kept in the functionally active oxidized and reduced states, respectively, by specific interactions with the cognate inner membrane oxidase DsbB and reductase DsbD, respectively. While early studies on Dsb proteins have offered a detailed mechanistic understanding of the classical pathways in *E. coli* K-12 (Zapun *et al.*, 1995, Bardwell *et al.*, 1993, Inaba *et al.*, 2006, Missiakas *et al.*, 1993, Martin *et al.*, 1993, Bardwell *et al.*, 1991, Stewart *et al.*, 1999, Kadokura *et al.*, 2004), much remains to be understood about the diversity in disulfide bond formation systems across bacteria (Heras *et al.*, 2009). It is now established that different bacteria can employ distinct Dsb machineries and encode, in addition to the classical DsbA/B and DsbC/D systems, multiple diverse periplasmic thiol/disulfide oxidoreductases (Heras *et al.*, 2009, Shouldice *et al.*, 2011, Landeta *et al.*, 2018). One such example is the suppressor of copper sensitivity (Scs) proteins (Gupta *et al.*, 1997).

Scs proteins were first discovered as the encoded products of a gene cluster associated with copper tolerance in *Salmonella enterica* serovar Typhimurium (Gupta *et al.*, 1997). The gene cluster encodes four proteins, ScsA, ScsB, ScsC, and ScsD. A putative copper-binding site and a thioredoxin-like motif containing the catalytic Cys-X-X-Cys site characteristic of Dsb proteins are present in most Scs proteins characterised to date (Petit *et al.*, 2022, Gupta *et al.*, 1997, Shepherd *et al.*, 2013, Furlong *et al.*, 2017, Cho *et al.*, 2012, Subedi *et al.*, 2021). Limited information on the biological function of ScsA and ScsD suggested a role in *S*. Typhimurium resistance to H_2_O_2_ and copper (López *et al.*, 2018). In comparison, the membrane-bound ScsB protein and soluble periplasmic ScsC protein have been characterized in three Enterobacteriaceae species to date: *S*. Typhimurium, *Proteus mirabilis*, and *Caulobacter crescentus* (Shepherd *et al.*, 2013, Furlong *et al.*, 2017, Cho *et al.*, 2012, Subedi *et al.*, 2021, Petit *et al.*, 2022). ScsB and ScsC constitute a redox pair that is functionally reminiscent of the *E. coli* DsbC and DsbD system (Shepherd *et al.*, 2013, Furlong *et al.*, 2017, Cho *et al.*, 2012, Subedi *et al.*, 2021, Petit *et al.*, 2022), in that ScsB catalytically reduces ScsC *in vivo* (Furlong *et al.*, 2017, Cho *et al.*, 2012). I*n vitro* evidence suggests ScsC scavenges toxic copper in the cells and delivers it to the copper-binding metallochaperone CueP, thereby reducing cellular toxicity in *S.* Typhimurium (Shepherd *et al.*, 2013, Subedi *et al.*, 2021, Petit *et al.*, 2022).

On the other hand, crystal structures of the ScsC homologues from *P. mirabilis* (PmScsC) and *C. crescentus* (CcScsC) revealed that their N-terminal regions oligomerize to form strikingly similar trimeric arrangements (Fig 1A-B) (Furlong *et al.*, 2017, Petit *et al.*, 2022). Nonetheless, the catalytic domains in each ScsC protomer resembles that of monomeric *E. coli* DsbA (Martin *et al.*, 1993) (Fig 1B-C). StScsC and CcScsC proteins were reported to function in their reduced state (Cho *et al.*, 2012, Subedi *et al.*, 2019). Specifically, the reduced thiols at the catalytic site of StScsC were directly implicated in copper-binding (Subedi *et al.*, 2021), with no significant disulfide isomerase (or oxidase) activity observed *in vitro* (Shepherd *et al.*, 2013, Subedi *et al.*, 2021). Conversely, the trimeric PmScsC and CcScsC proteins had disulfide isomerase activity and lacked dithiol oxidase activity (Furlong *et al.*, 2017, Cho *et al.*, 2012).

**Fig. 1.**
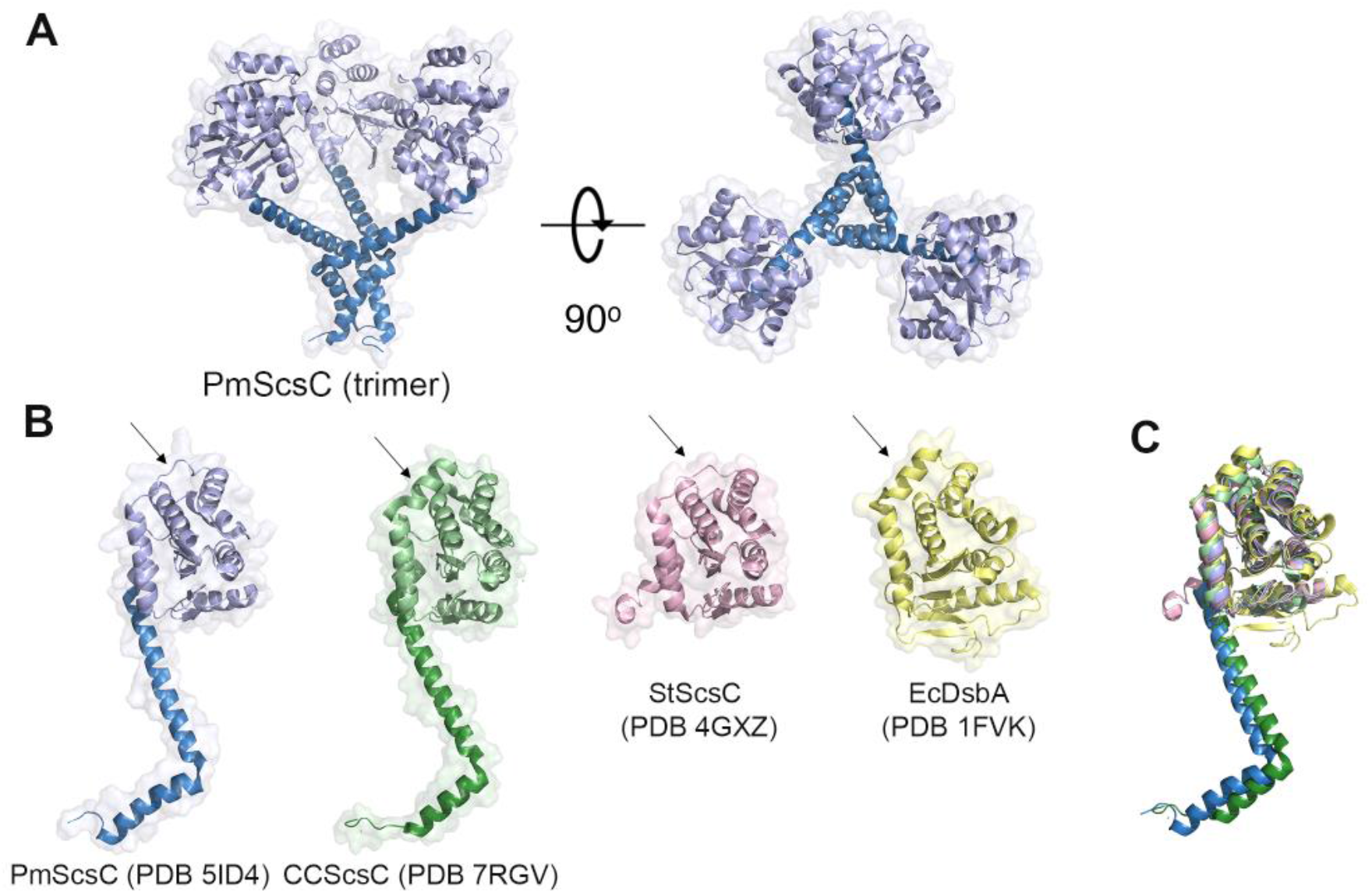
Structural similarity between ScsC homologues and DsbA. A, The trimeric architecture of PmScsC protein, PDB 5id4 (Furlong *et al.*, 2017). Trimerization and catalytic domains are displayed in blue and slate, respectively. B, Protomer structures of PmScsC (blue/slate; PDB 5id4 (Furlong *et al.*, 2017)), CcScsC (green; PDB 7rgv (Petit *et al.*, 2022), this protein is a biological trimer), StScsC (pink; PDB 4gxz (Shepherd *et al.*, 2013), monomeric) and EcDsbA (yellow; PDB 1fvk (Guddat *et al.*, 1998), monomeric). The α-helix/loop differences in the globular domains are indicated with an arrow. C. Structural superposition of PmScsC, CcScsC, StScsC and EcDsbA.

Considerable progress has been made over the past decades in the genetics, mechanism, and structural understanding of thiol-disulfide oxidoreductases. Nonetheless, what governs their biochemical fate to exhibit either dithiol oxidase or disulfide reductase activity remains a major open question. Collectively, evidence from individual studies suggests a role for thioredoxin fold oligomerization in determining specific thiol oxidoreductase catalytic activity. For example, globular periplasmic monomeric oxidoreductases like EcDsbA are dithiol oxidases, whereas the V-shaped periplasmic homodimeric Dsb proteins EcDsbC and EcDsbG exhibit disulfide isomerase activity (Hiniker *et al.*, 2007, Santos-Martin *et al.*, 2021, Zapun *et al.*, 1995, Banaszak *et al.*, 2004, Heras *et al.*, 2004). Furthermore, when EcDsbC was engineered to be monomeric, it acquired oxidase activity at the cost of its original disulfide isomerase activity (Zhao *et al.*,2003, Bader *et al.*, 2001). Conversely, when EcDsbA was dimerised, the protein gained disulfide isomerase activity, while its oxidase activity was impacted (Segatori *et al.*, 2004).

The three structurally characterised ScsC proteins, StScsC, CcScsC, and PmScsC, adopt distinct oligomeric states and exhibit distinct oxidoreductase activities *in vitro*, despite their catalytic domains sharing a striking structural resemblance with monomeric EcDsbA (Fig. 1) (Furlong *et al.*, 2017, Petit *et al.*, 2022, Shepherd *et al.*, 2013, Cho *et al.*, 2012, Subedi *et al.*, 2021). Furthermore, similar to other Dsb proteins, upon the loss of the N-terminal trimeric module, monomeric PmScsC protein (PmScsCΔN) was reported to lose disulfide reductase activity and gain dithiol oxidase activity *in vitro* (Furlong *et al.*, 2017). Here, we set out to investigate in a cellular context how the quaternary arrangement of thioredoxin-like oxidoreductases influences ScsC function. We used the set of three previously described ScsC homologues (monomeric StScsC and trimeric PmScsC and CcScsC) and the shared genetic background of the model organism *E. coli* K-12.

Herein, we report that all three ScsC proteins possess both dithiol oxidase and disulfide reductase cellular activity, irrespective of their oligomeric state (monomeric or trimeric). Intriguingly, both StScsC and CcScsC but not PmScsC required the EcDsbD electron donor for effective reductase activity in *E. coli* K-12 in the absence of a cognate ScsB partner. Contrastingly, dithiol oxidase activity of ScsC proteins was restricted by the lack of a compatible redox partner in *E. coli* K-12 and was only observed *in vivo* under conditions of environmental oxidation. An engineered monomeric form of PmScsC was the exception, as it could use EcDsbB (the partner of EcDsbA) to efficiently catalyse dithiol oxidation in cells. This study systematically explored the contribution of protein oligomerisation in ScsC cellular activity, revealing that it could serve as one strategy by which bacteria can limit redox protein-protein interactions in the cell to maintain oxidoreductase activities that are compatible with the different and diverse Dsb systems they encode.

## MATERIAL AND METHODS

### Bacterial strains, plasmids, and growth conditions

The strains and plasmids used in this study are described in Tables 1 and 2, respectively. Strains were grown at 37°C in LB-Lennox (5 g L^-1^ NaCl, 5 g L^-1^ yeast extract, 10 g L^-1^ tryptone) or on agar plates (LB with 15 g L^-1^ agar) unless otherwise indicated. Wherever appropriate, antibiotics were added at the following concentrations: ampicillin (100 *μ*g mL^-1^), kanamycin (50 *μ*g mL^-1^), and chloramphenicol (17 *μ*g mL^-1^).

**Table 1.**
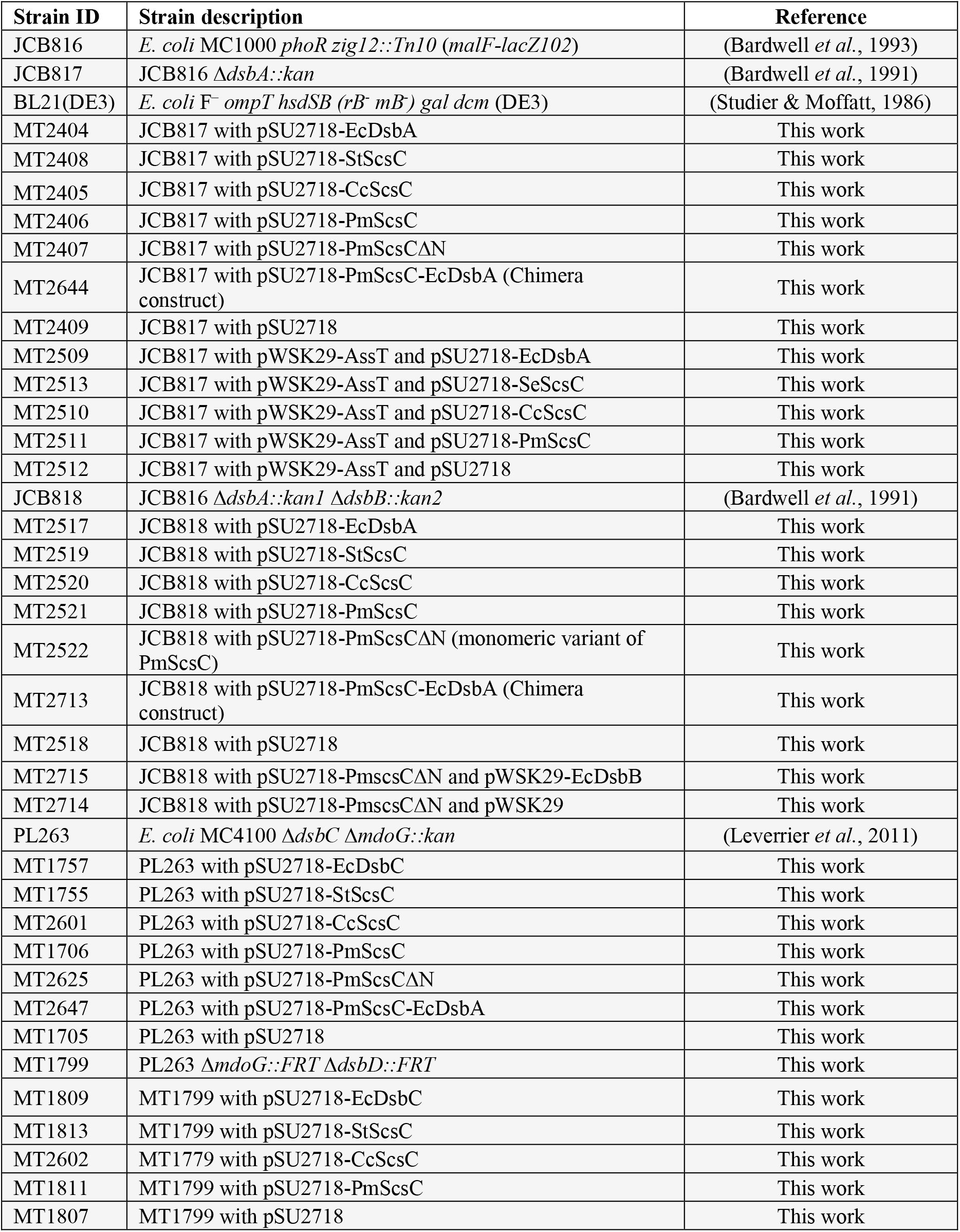
Bacterial strains used in this study

**Table 2.**
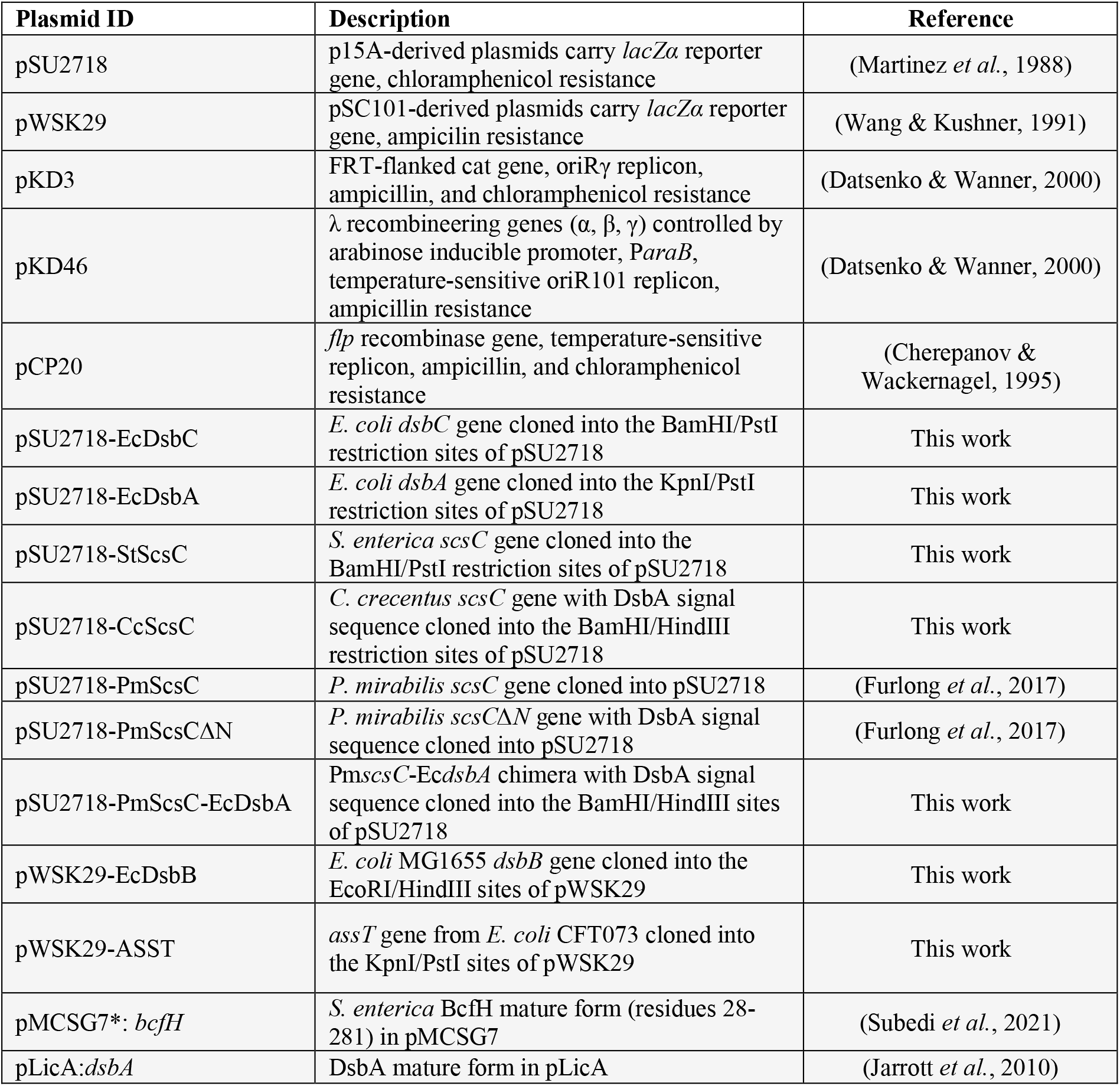
Plasmids used in this study.

### Bacterial genetics methods

PmScsC-EcDsbA chimera was generated by Gibson assembly. The catalytic domains of *E. coli* DsbA (EcDsbA^Cat^) was amplified by polymerase chain reaction (PCR) using the primers described in Table S1. The previously reported pMCSG7-PmScsC construct (Furlong *et al.*, 2017) was used as the template for PCR amplification of the N-terminal trimerization domain of PmScsC using the primers shown in Table S1. The gene encoding the PmScsC-EcDsbA chimera was amplified with NcoI-Pm*scsC* (Δss) Forward, and HindIII-EcDsbA Reverse (Table S1) from the expression construct into pSC108 (Cho *et al.*, 2012) so that the chimera carries an inframe DsbA signal sequence to aid the translocation of the protein. The resultant DsbAss--Δss(PmscsC)-PmscsC-EcDsbA chimera was then subcloned into the BamHI/HindIII sites of pSU2718 (Martinez *et al.*, 1988). The *C. crescentus scsC* gene (with *E. coli* DsbA signal sequence) was amplified from pSC108 (DsbAss-CcScsC) (Cho *et al.*, 2012) with *BamHI-dsbA* Forward and HindIII-Cc*scsC* Reverse (see Table S1) and cloned into pSU2718 (Martinez *et al.*, 1988). The *S. enterica scsC* gene was subcloned from pWSK29-StScsC (Shepherd *et al.*, 2013) into the BamHI/PstI restriction sites of pSU2718 (Martinez *et al.*, 1988). Plasmids and knockout mutations in this work were constructed using a modified version of the Datsenko and Wanner method (Datsenko & Wanner, 2000, Hong & Reeves, 2014). Wherever appropriate, the selection cassettes used for gene knockouts were cleaved by Flp recombinase (Cherepanov & Wackernagel, 1995). Polyethylene-glycol-induced transformations were used to transfer plasmids into the testing strains constructed for this work (Klebe *et al.*, 1983). P1 Phage cultivation, purification and enrichment were performed as previously described (Thomason *et al.*, 2007). The transfer of genetic traits between strains was performed by P1 transductions using the Thomason *et al.* (2007) procedure. For the oligonucleotides used in cloning and gene replacements, see Supplemental material (Table S1).

### Swimming motility assay

Bacterial swimming motility was measured as described previously with modified conditions (Verderosa *et al.*, 2021). Briefly, overnight 24 hr static cultures were normalized to OD_600_ 1.0 (~ 1 × 10^9^ CFU/mL) and inoculated onto 0.25% w/v LB-Lennox agar (Gibco^™^ Bacteriological agar) containing IPTG (500 *μ*M) and chloramphenicol (17 *μ*g mL^-1^) that had been left to set and then dried overnight at room temperature (21 °C). Swimming motility was then assessed by tracking OD_600_ in a CLARIOstar plate reader (BMG, Australia) at 37 °C over 20 hrs.

### ASST sulfotransferase assay

Sulfotransferase activity was measured as previously described (Verderosa *et al.*, 2021) with modifications. Briefly, bacterial strains were cultured in LB-Lennox media at 37 °C overnight with 200 rpm aeration in the presence of IPTG (500 *μ*M), ampicillin (100 *μ*g mL^-1^) and chloramphenicol (17 *μ*g mL^-1^). The cultures were washed once in 1× PBS, normalized to OD_600_ 0.8 (~ 7 × 10^8^ CFU mL^-1^) and transferred 100 *μ*L per well into a Costar^®^ 96-well plate. Cultures were mixed with an equal volume of 1× PBS solution containing 1 mM potassium 4-methylumbelliferyl sulfate (SIGMA ALDRICH, Castle Hill, Australia) and 2 mM phenol (SIGMA ALDRICH, Australia). Sulfotransferase activity was monitored in a CLARIOstar plate reader (BMG, Australia) over a 90 mins period by measuring the fluorescence emitted at 450–480 nm (excitation wavelength at 360–380 nm).

### Mucoidal colony morphology assay to track colanic acid production

Disulfide isomerase activity by ScsC proteins was determined phenotypically by enhanced production of colanic acid in *E. coli* K-12 similar to previously described (Leverrier *et al.*, 2011). Briefly, single bacterial colonies were patched onto 1.5% solid agar M9 minimal media (2 μM MgSO_4_, 100 nM CaCl_2_, 0.4% v/v glycerol and 0.1% w/v casamino acid) containing IPTG (500 *μ*M), and wherever appropriate, relevant antibiotics were added to maintain plasmids. The Cold Spring Harbor Protocol formula for M9 salts was used in this study (10× stock, per L, 70 g Na_2_HPO_4_•7H_2_O, 30 g KH_2_PO_4_, 5 g NaCl, 10 g NH_4_Cl). Plates were incubated at 29 °C for 48 hrs and macroscopically visualised for mucoidal colony morphology. Images were taken using a ChemiDoc MP Imaging System (Bio-Rad).

### Purification of PmScsC-EcDsbA chimera

Large scale expression of PmScsC-EcDsbA chimera was carried out by autoinduction in *E. coli* BL21(DE3) (Studier, 2005), with incubation at 37°C for 4 h with 180 rev min^-1^ agitation, followed by incubation at 30°C for 18 hr with 180 rev min^-1^ agitation. PmScsC-EcDsbA was initially purified by immobilized metal affinity chromatography (IMAC) using a 5 mL HisTrap Fast Flow column (GE Healthcare) followed by cleavage of the N-terminal His6-tag with TEV protease and reverse IMAC. The chimera was purified to homogeneity by size exclusion chromatography.

### Biophysical characterisation

Analysis of protein secondary structure of PmScsC-EcDsbA was conducted by circular dichroism (CD) spectroscopy with an AVIV Model 410SF circular dichroism spectrophotometer (Aviv Biomedical, Inc) using a 1 mm quartz cuvette. 200 μg ml^-1^ of oxidised or reduced protein was prepared in 20 mM sodium phosphate, 0.1 mM EDTA, pH 7, with reduced protein samples supplemented with 0.75 mM DTT. Wavelength scans at 196–250 nm were conducted at 20°C and the CD signal from each protein scan were buffer subtracted and converted to molar ellipticity (Δε (M^-1^ cm^-1^). Analytical size exclusion chromatography was performed with 5 mg of EcDsbA, BcfH (a trimeric Dsb protein) or PmScsC-EcDsbA [1.25 mg ml^-1^] using a Superdex^®^ 200 Increase 10/300 GL column equilibrated with 25mM HEPES pH 6.5, 150 mM NaCl. Elution profiles were determined by measuring absorbance at 280 nm. Sedimentation velocity analytical ultracentrifugation (SV-AUC) was used to confirm the oligomerization state of the chimera PmScsC-EcDsbA. SV-AUC was performed at 20°C using a Beckman Optima XL-A analytical ultracentrifuge using a conventional double sector quartz cell mounted in an 8-hole An-50 Ti rotor. SV-AUC experiments were conducted using 380 μL of 2 mg ml^-1^ protein and 400 μL of reference solution consisting of 25 mM HEPES, 150 mM NaCl, pH 7.0. SV-AUC was run at 40,000 rpm and scans were collected in a continuous mode at 285 nm. The partial specific volume, solvent density and viscosity were computed using SEDNTERP (Lebowitz *et al.*, 2002) and data was fitted using SEDFIT (Schuck *et al.*, 2002).

## RESULTS

### ScsC homologues can restore loss of disulfide oxidase and isomerase pathways in E. coli

To assess the different ScsC homologues for disulfide isomerase activity *in vivo* we employed an established *E. coli* assay that monitors colanic acid production (Leverrier *et al.*, 2011). The previously described *E. coli* K-12 strain MC4100 Δ*dsbC* Δ*mdoG* (PL263) (Leverrier *et al.*, 2011) was transformed with pSU2718-based expression plasmids containing *scsC* genes from *S.* Typhimurium, *P. mirabilis*, and *C. crescentus* or Ec*dsbC* as the positive control. The deletion of *mdoG* in this *E. coli* strain triggers cell envelope stress, thereby activating RcsF-dependent colanic acid biosynthesis that leads to a visually distinctive mucoidal colony morphology on agar plates (Majdalani *et al.*, 2005, Castanié-Cornet *et al.*,2006). This assay monitors *in vivo* disulfide isomerase activity, as the outer membrane RcsF protein contains two pairs of disulfide bonds formed between nonconsecutive Cys residues; thus, correct RcsF folding and subsequent activation of the Rcs signalling cascade that leads to a mucoidal phenotype, requires the presence of an active disulfide isomerase, such as EcDsbC (Leverrier *et al.*, 2011).

MC4100 Δ*dsbC* Δ*mdoG* strains containing *scsC* expression plasmids were assessed for colanic acid production on minimal M9 media (Fig. 2A). Colanic acid biosynthesis (resulting in mucoid colony appearance) was restored in the presence of the native isomerase EcDsbC, as expected (Leverrier *et al.*, 2011) (positive control strain MC4100 Δ*dsbC* Δ*mdoG*/pEc*dsbC* (Fig 2A)). Expression of CcScsC also complemented the disulfide isomerase activity defect in MC4100 Δ*dsbC* Δ*mdoG* (Fig. 2A), consistent with a previous report by (Cho *et al.*, 2012). Complementation with StScsC led to mucoidal colony morphology, evidencing isomerase activity for this protein for the first time. The trimeric PmScsC previously reported to possess isomerisation activity *in vitro* (Furlong *et al.*, 2017), was now confirmed to function as an isomerase *in vivo* (Fig. 2A).

**Fig. 2.**
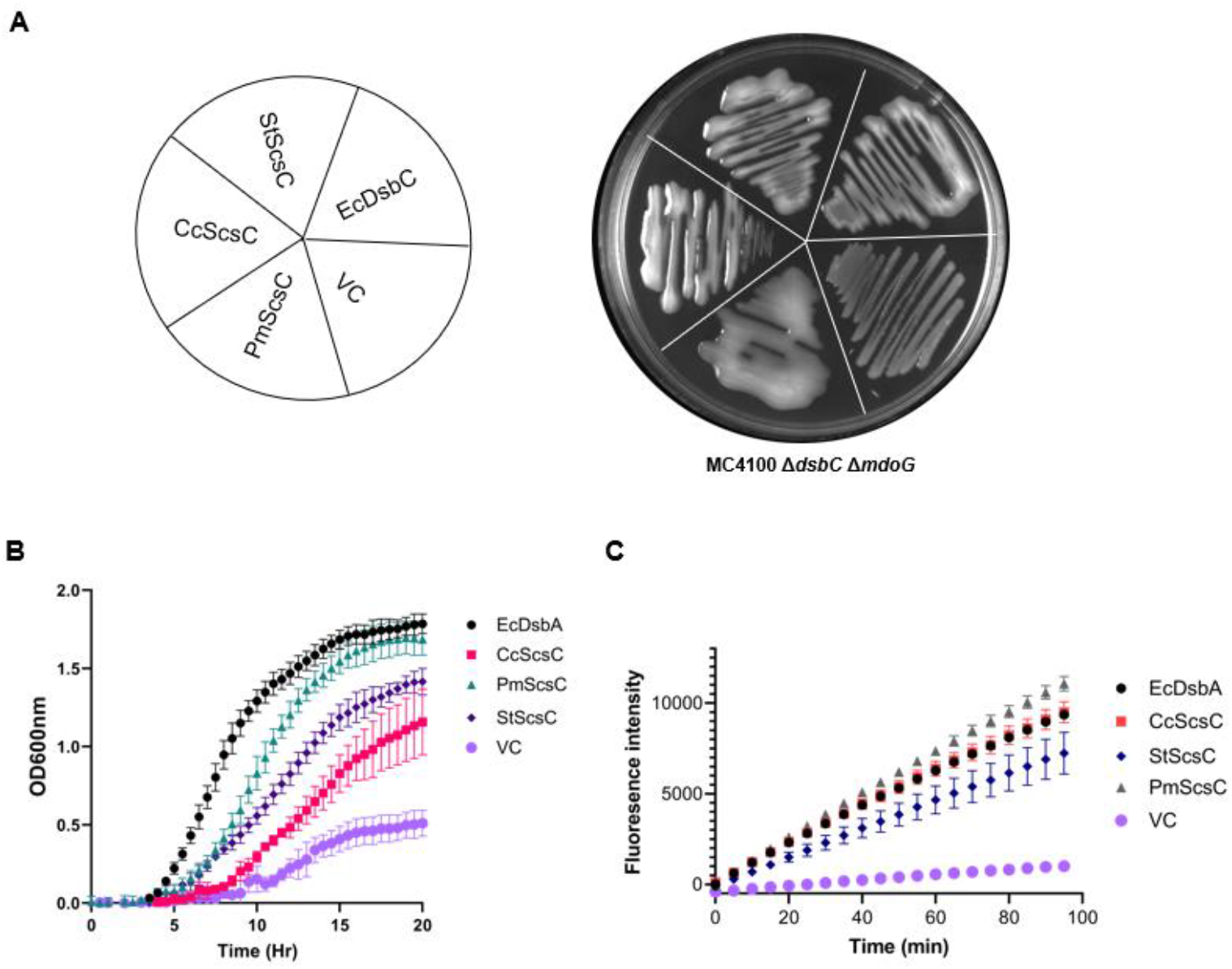
ScsC proteins display both disulfide isomerase and oxidase activity in *E. coli.* A, Complementation of RcsF folding and restoration of mucoidal colony morphology in strain MC4100 Δ*dsbC* Δ*mdoG* by disulfide isomerase EcDsbC (positive control) and ScsC proteins expressed from the same plasmid vector, or empty vector control (VC). Plate image (representative of three independent repeats) shows complemented strains grown on solid M9 minimal media following incubation at 29°C for 48 hrs. B, Complementation of FlgI folding and restoration of flagella-mediated swimming motility in strain JCB816 Δ*dsbA* by EcDsbA and ScsC proteins, or empty VC. Strain motility zones on semi-solid media were measured as optical density (OD_600nm_) every five minutes over 20 hours of continuous incubation at 37°C. C, Complementation of ASST folding and restoration of cellular sulfotransferase activity in strain JCB816 Δ*dsbA* pAssT by EcDsbA and ScsC proteins, or empty VC. ASST activity was monitored by fluorogenic substrate intensity measured every 5 minutes over 95 minutes; Data in panels B and C show mean values derived from biological triplicates with error bars representing the standard deviation (SD).

To examine if ScsC proteins have oxidase activity *in vivo*, we employed the *E. coli* swimming motility assay, as introduction of disulfide bonds in FlgI is required for swimming motility in *E. coli* (Dailey & Berg, 1993, Hizukuri *et al.*, 2006). FlgI is a component of the *E. coli* flagellum motility organelle that contains an intramolecular disulfide bond catalysed by EcDsbA. The non-motile JCB816 Δ*dsbA* strain was transformed with *scsC* expression plasmids and compared to EcDsbA to restore swimming motility. All three of the ScsC proteins restored motility in the *dsbA* null strain (Fig. 2B), with PmScsC restoring *E. coli* motility to similar levels as EcDsbA (positive control). These findings were confirmed in a separate assay that monitors *in vivo* oxidative folding of a different established DsbA substrate, the periplasmic enzyme arylsulfate sulfotransferase (ASST). ASST contains a disulfide bond required for biochemical activity (Kwon & Choi, 2005) and is found in several bacterial species, including uropathogenic *E. coli* (Grimshaw *et al.*, 2008, Totsika *et al.*, 2009). A plasmid containing the *assT* gene (pASST) was transformed into the same set of *E. coli* JCB816 Δ*dsbA* strains used in the motility assays, and ASST oxidative folding by ScsC proteins was compared to the EcDsbA positive control. All ScsC homologues restored ASST activity in the *dsbA* null mutant at comparable or slightly reduced levels to EcDsbA, confirming ScsC oxidase activity *in vivo* (Fig. 2C). Collectively, these results demonstrate that despite previous reports of ScsC enzymes acting predominantly as isomerases *in vitro*, *in vivo* they can act both as isomerases and oxidases and participate in mainstream oxidative protein folding pathways present in *E. coli* K-12.

### DsbD but not DsbB serves as a heterologous redox partner for ScsC in E. coli

To further investigate the participation of ScsC proteins in *E. coli* oxidative protein folding pathways, we examined if *in vivo* ScsC activity required DsbB and DsbD, which are the respective redox partners of DsbA and DsbC in *E. coli.* Interestingly, deletion of *dsbB* in the *E. coli* strains assessed for swimming motility did not alter the original strain phenotype, indicating that ScsC expression could alone restore *E. coli* motility in the absence of EcDsbA and EcDsbB (Fig. 3A and 3B). This was, however also observed with EcDsbA, which under the test conditions, could fully complement motility despite loss of EcDsbB (Fig. 3A and 3B). Full restoration of motility in a Δ*dsbB* strain was previously reported to occur with exogenously supplemented oxidants (Dailey & Berg, 1993) presumably through environmental oxidation of EcDsbA. For this reason, we repeated motility assays in minimal M9 media. Under the new test conditions, strains remained immotile (Fig. 3C), confirming that environmental oxidation of EcDsbA, as well as of ScsC proteins, was responsible for the motility restoration observed in rich media. In contrast, in minimal media, the motility of the JCB816 Δ*dsbA* (DsbB+) strain was only fully restored by complementation with EcDsbA, while ScsC expressing strains remained immotile, with the exception of the strain expressing PmScsC, which showed marginal complementation of motility under the test conditions (Fig. 3D). We conclude that ScsC proteins can be active as oxidases in *E. coli* K-12 and fold different DsbA substrates under oxidant supplementation conditions, as their oxidase activity is not maintained by DsbB *in vivo*.

**Fig. 3.**
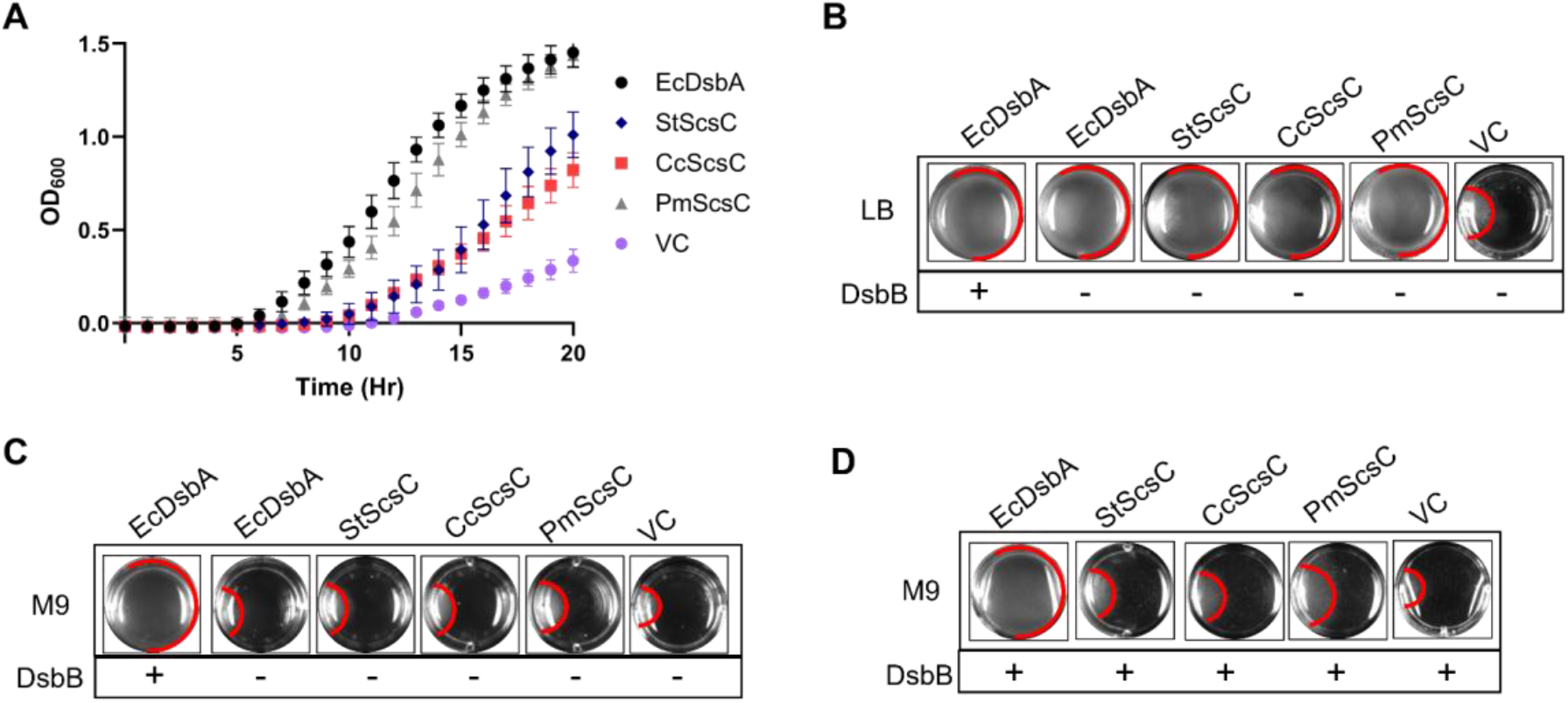
Dithiol oxidase activity of ScsC proteins in *E. coli* is dependent on exogenous oxidants. A and B. Expression of EcDsbA and ScsC proteins but not empty vector control (VC) restores swimming motility in JCB816 Δ*dsbA* Δ*dsbB* in rich media (panel A shows motility kinetics monitored by OD_600_ measured every 5 minutes over 20 hours and panel B shows motility plate image taken at 20 hrs with swimming zone edge marked in red); C. Expression of ScsC proteins does not restore swimming motility in JCB816 Δ*dsbA* Δ*dsbB* in minimal media (plate image taken at 20 hours); D. Expression of EcDsbA but not ScsC proteins fully restores swimming motility in JCB816 Δ*dsbA* (DsbB+) in minimal media (plate image taken at 20 hours). Data plotted in A are mean values from biological triplicates with SD values shown as error bars. The images shown in panels B-D are representative of at least three biological replicates.

The lack of DsbB dependency for oxidase activity by the ScsC proteins *in vivo* prompted us to investigate their requirement for DsbD as a redox partner for isomerase activity next. For this, we constructed a *dsbD* null mutation in MC4100 Δ*dsbC* Δ*mdoG*, the reporter strain for isomerase activity. When the Δ*dsbC* Δ*mdoG* knockout strain was complemented with EcDsbC alone, the hallmark mucoidal morphology induced by RcsF folding was observed (Fig. 4). This mucoidal morphology was reduced in the MC4100 Δ*mdoG* Δ*dsbD* Δ*dsbC* strain confirming DsbC dependence on DsbD for isomerase activity. Similarly, expression of StScsC and CcScsC in a strain lacking DsbD and DsbC showed reduced colanic acid production compared to an isogenic strain with DsbD (Fig. 4). In contrast, strains expressing PmScsC retained high colanic acid production irrespective of DsbD presence or absence (Fig. 4). These results suggest that, like EcDsbC, efficient disulfide isomerase activity by StScsC and CcScsC, but not by PmScsC, in *E. coli* requires DsbD.

**Fig. 4.**
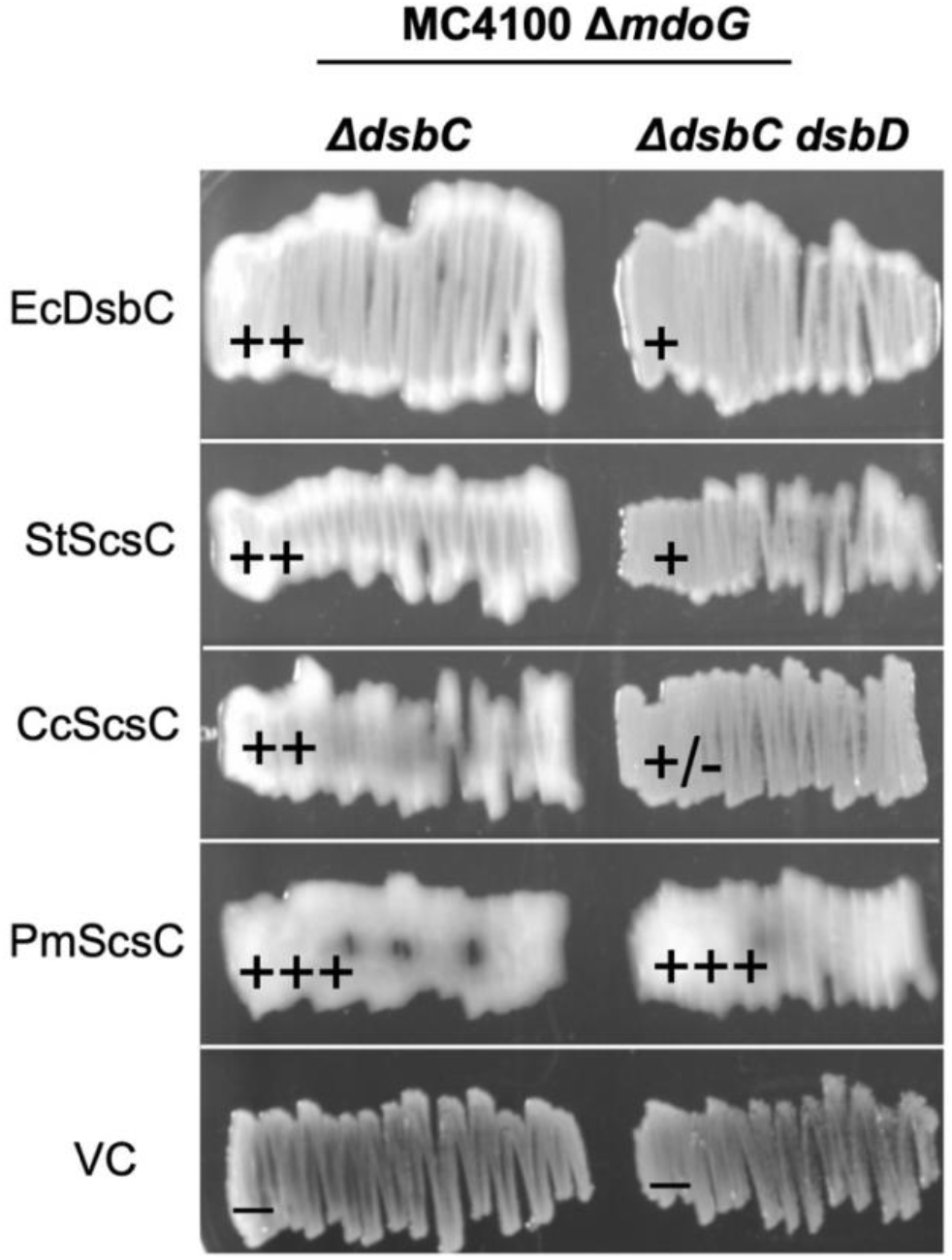
Requirement for DsbD in maintaining ScsC disulfide isomerase activity in *E. coli.*Restoration of RcsF-dependent colanic acid production and mucoidal colony morphology in strains MC4100 Δ*mdoG* Δ*dsbC* (left) and MC4100 Δ*mdoG* Δ*dsbC* Δ*dsbD* (right) by EcDsbC and ScsC proteins expressed from the same plasmid vector, or empty vector control (VC). The plate image shows complemented strains grown on solid minimal media following incubation at 29°C for 48 hrs (plate image is representative of at least three biological replicates). Intensity of mucoidal appearance is scored as strong ‘+++’ or ‘++’, weak ‘+’ or ‘+/-’, or none ‘-’.

### Monomeric PmScsC requires DsbB to function as a dithiol oxidase in E. coli

The unique activity of PmScsC as an efficient DsbD-independent isomerase, combined with a strong capacity to fold DsbA substrates upon becoming environmentally oxidised, prompted us to investigate this intriguing disulfide catalyst further. Its quaternary structure consists of three protomers that are tethered together into a trimer (Fig. 1) (Furlong *et al.*, 2017). Trimerization occurs through the oligomerization of a modular N-terminal trimerization domain (Furlong *et al.*, 2017). Full-length PmScsC failed to oxidize dithiol peptides *in vitro* (Furlong *et al.*, 2017), but upon removing the trimerization domain (PmScsCΔN) the protein adopted a monomeric structure that acquired oxidase activity at the expense of isomerase activity *in vitro* (Furlong *et al.*, 2019, Furlong *et al.*, 2017). We hypothesized that monomeric PmScsCΔN also attains the ability to interact with EcDsbB and function as an efficient oxidase in *E. coli.* To test this, we compared motility restoration in strain JCB816 Δ*dsbA* by PmScsC and PmScsCΔN in minimal media. Under these conditions, only the monomeric PmScsCΔN restored bacterial motility to EcDsbA levels (Fig. 5A). Motility restoration by PmScsCΔN was dependent on DsbB, similar to EcDsbA, as the expression of PmScsCΔN in JCB816 Δ*dsbA* Δ*dsbB* failed to restore bacterial motility, unless the strain was complemented with a plasmid expressing EcDsbB (Fig. 5B). Moreover, PmScsCΔN lost isomerase activity in *E. coli* in contrast to trimeric PmScsC (Fig. 5C). Together, our results confirm that monomeric (PmScsCΔN) but not trimeric PmScsC is an effective thiol oxidase that forms a functional redox pair with DsbB catalysing oxidative protein folding in the *E. coli* periplasm.

**Fig. 5.**
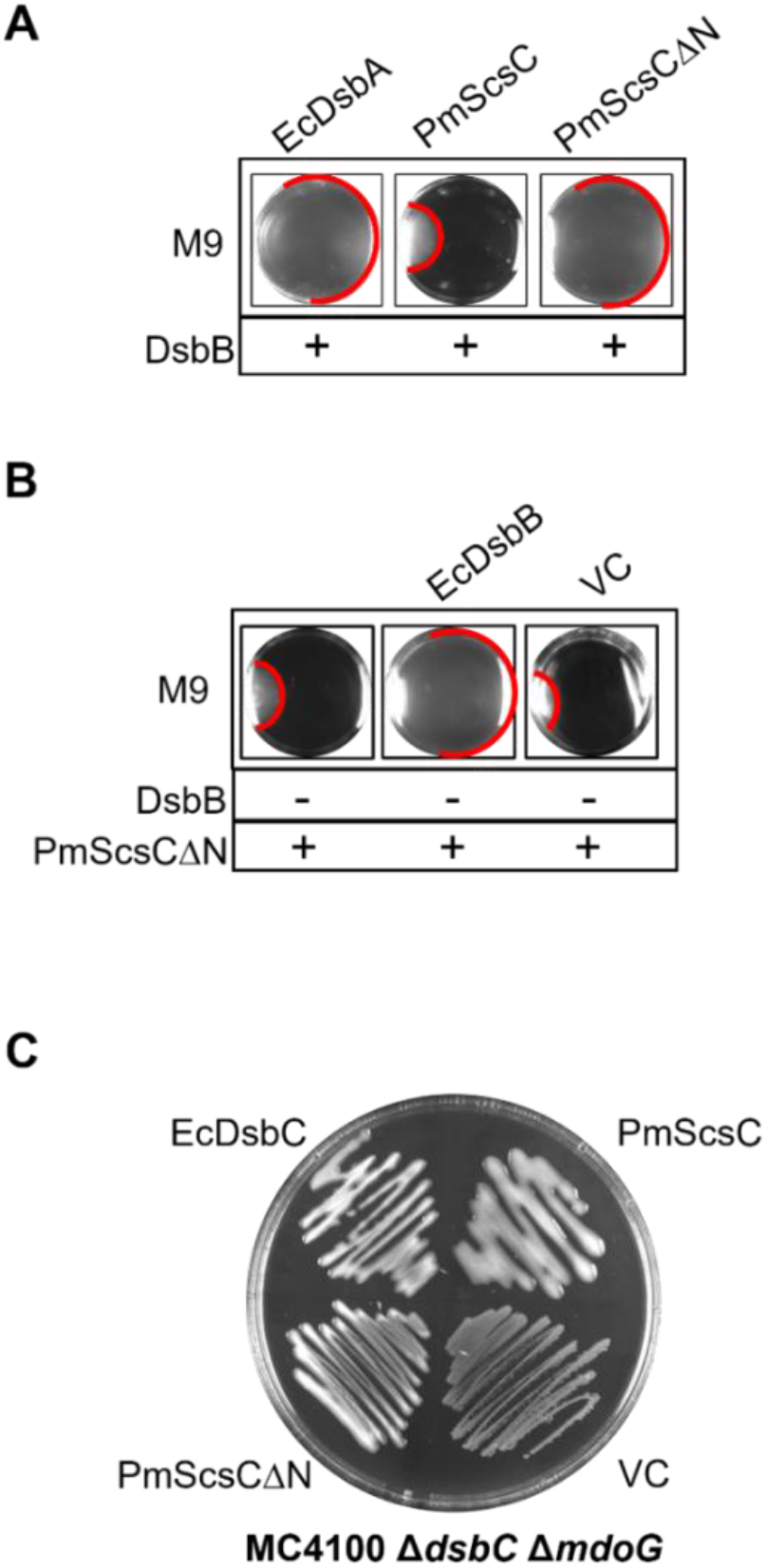
The oligomeric state of PmScsC dictates its disulfide oxidoreductase activity and redox protein interactions in *E. coli.* A, Restoration of JCB816 Δ*dsbA* (DsbB+) motility by disulfide oxidase EcDsbA (positive control), PmScsC (trimeric) and PmScsCΔN (monomeric) in minimal media; B, Restoration of JCB816 Δ*dsbA* Δ*dsbB* (DsbB-) motility by PmScsCΔN alone, or PmScsCΔN and plasmid-borne EcDsbB or empty VC; C, Restoration of RcsF-dependent colanic acid production in strain MC4100 Δ*mdoG* Δ*dsbC* by disulfide isomerase EcDsbC (positive control), PmScsC (trimeric), PmScsCΔN (monomeric) or empty VC. All images are representative of at least three biological replicates.

### Trimeric EcDsbA retains disulfide oxidase activity without gain of isomerase activity in E. coli

The change of catalytic activity observed in PmScsC from a disulfide isomerase to an oxidase purely by altering its oligomeric state, prompted us to investigate whether modulation of catalytic activity could be engineered into the prototypical monomeric *E. coli* thiol oxidase DsbA by oligomerization. For this, we constructed a PmScsC-EcDsbA chimeric protein that consists of the modular N-terminal PmScsC linker domain preceding the mature EcDsbA sequence. Over-expression of PmScsC-EcDsbA was carried out in BL21(DE3) and purified to homogeneity using IMAC, followed by TEV cleavage, reverse IMAC and size exclusion chromatography (SEC) (Fig 6A). The secondary structure of PmScsC-EcDsbA in both oxidised and reduced state was assessed by CD spectroscopy, which showed a predominantly α-helical secondary structure as reflected by the double minima at 209 and 222 nm, with the former being more significant, which is expected for folded DsbA-like proteins (Fig 6B). The trimeric oligomerisation state in solution of the PmScsC-EcDsbA chimera was analysed by size exclusion chromatography (SEC) and SV-AUC. SEC analysis showed that PmScsC-EcDsbA elutes at a similar elution volume as previously described trimeric DsbA-like proteins [PmScsC (molecular weight 74.3 kDa (Furlong *et al.*, 2017)) and BcfH (molecular weight 82.8 kDa, (Subedi *et al.*, 2021))] and earlier than monomeric EcDsbA (23.1 kDa), suggesting that the chimera oligomerises in solution (Fig 6C). Sedimentation velocity analytical ultracentrifugation (SV-AUC) was performed to confirm the molecular weight of PmScsC-EcDsbA. The results indicate that this chimera has a molecular weight of 79 kDa (Fig 6D), approximately 3-times the molecular weight of monomeric PmScsC-EcDsbA (27 kDa) and consistent with formation of a trimer. The AUC profile shows that the chimera displays a slightly broader peak than EcDsbA, which may reflect the presence of small amounts of a second species that migrates at slightly different rate. Altogether, these results indicate that the addition of the N-terminal trimerization of EcDsbA resulted in a trimeric PmScsC-EcDsbA chimera.

**Fig. 6.**
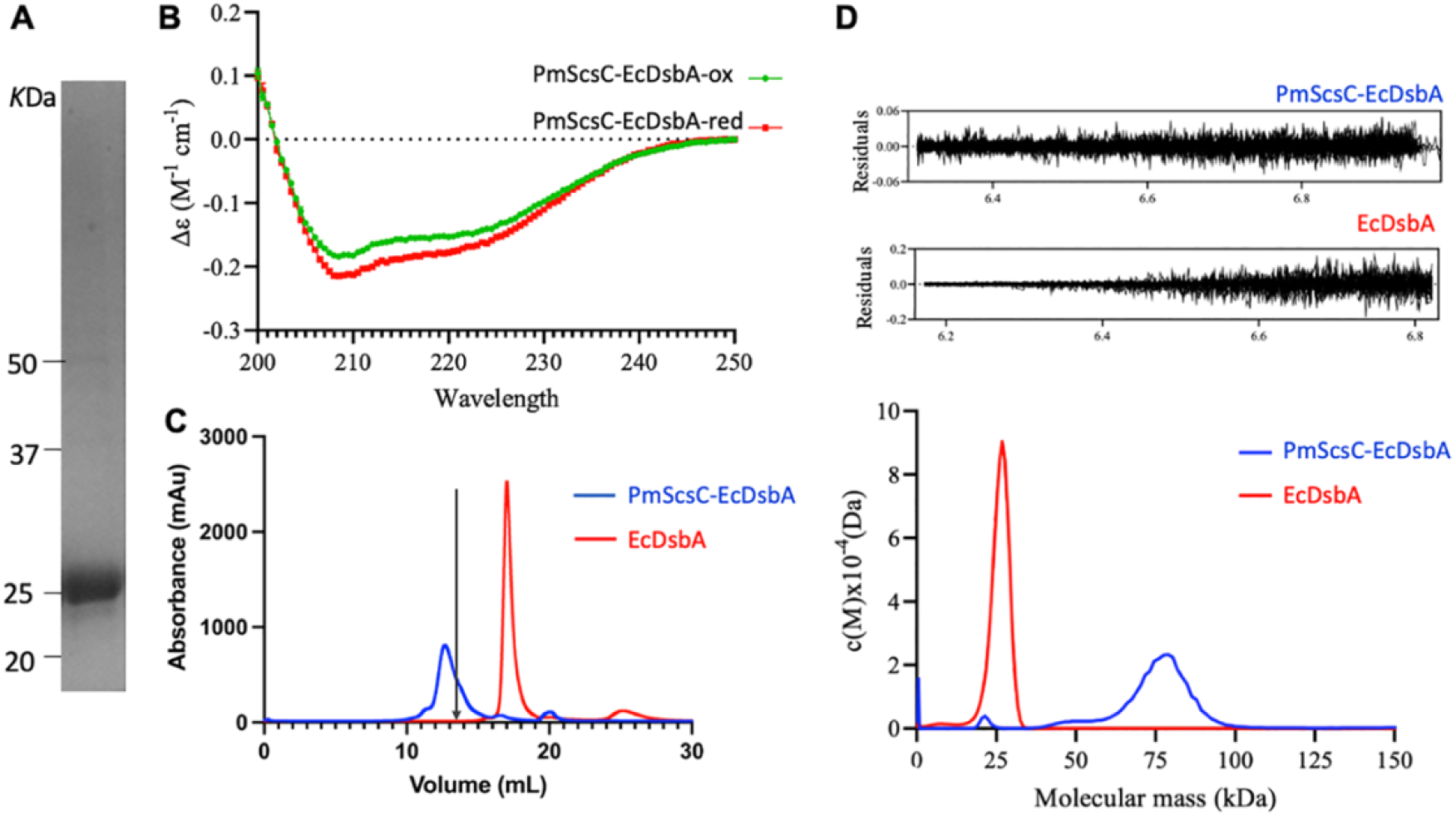
PmScsC-EcDsbA chimera production and biophysical characterisation. A. SDS-PAGE analysis of purified PmScsC-EcDsbA (monomer molecular weight 27 kDa). B. Circular dichroism spectra of oxidised and reduced PmScsC-EcDsbA. CD measurements were at 20°C and plotted as molar ellipticity Δε (M^-1^ cm^-1^) as a function of the wavelength (200 – 250 nm). C. Size exclusion chromatography elution profiles PmScsC-EcDsbA (blue) and monomeric EcDsbA (red). Black arrow indicates the elution volume for trimeric DsbA-like proteins (Furlong *et al.*, 2017, Subedi *et al.*, 2021). D. Sedimentation velocity analytical ultracentrifugation analysis displaying the continuous mass [c(M)] distribution as a function of molecular mass (kDa) of PmScsC-EcDsbA and EcDsbA at 2 mg ml^-1^ with residuals for *c*(M) of PmScsC-EcDsbA and EcDsbA displayed above.

To assess the activity of trimeric EcDsbA *in vivo*, we expressed the PmScsC-EcDsbA chimera in *E. coli* and assessed its capacity to restore swimming motility and colanic acid production in relevant strains. Surprisingly, full motility restoration of strain JCB816 Δ*dsbA* was mediated by the trimeric DsbA chimera, at similar levels to wild-type monomeric EcDsbA (positive control) (Fig. 7A). Like monomeric EcDsbA, the trimeric protein failed to restore colanic acid production in MC4100 Δ*dsbC* Δ*mdoG* (Fig. 7B). Collectively our findings demonstrate that trimerization of EcDsbA does not affect its strong thiol oxidation activity or impart it with an additional isomerisation function in the cell.

**Fig. 7.**
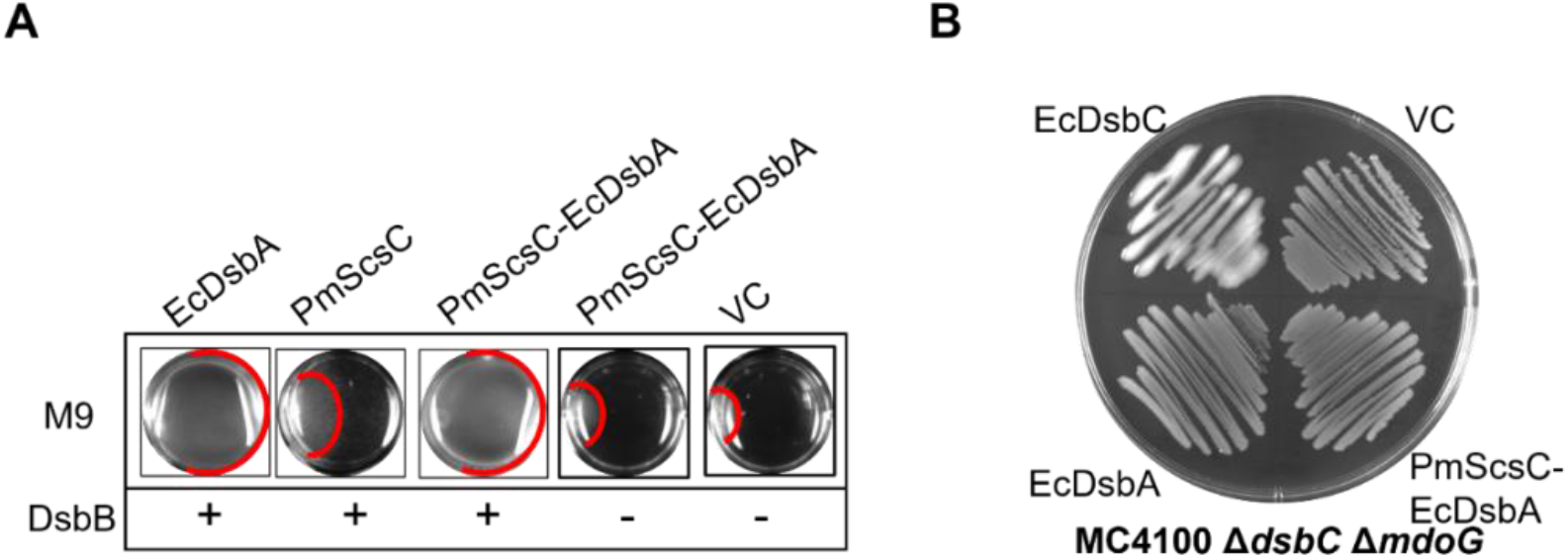
Trimerization of EcDsbA does not affect its disulfide oxidoreductase function in *E. coli.*A, Restoration of swimming motility in minimal media for strain JCB816 Δ*dsbA* (DsbB+) expressing EcDsbA (monomeric), PmScsC (trimeric) and PmScsC-EcDsbA chimera (trimeric) and isogenic *dsbB* mutant JCB816 Δ*dsbA* Δ*dsbB* (DsbB-) complemented with the PmScsC-EcDsbA chimera or empty VC; B, Restoration of RcsF-dependent colanic acid production in strain MC4100 Δ*mdoG* Δ*dsbC* by disulfide isomerase EcDsbC (positive control), EcDsbA (monomeric), PmScsC-EcDsbA chimera (trimeric) or empty VC.

## DISCUSSION

Disulfide bond catalysis by bacterial Dsb proteins governs the oxidative folding of several extra-cytoplasmic proteins in the periplasmic space (Heras *et al.*, 2009, Shouldice *et al.*, 2011, Landeta *et al.*, 2018, Dutton *et al.*, 2008). This oxidative protein folding process plays a crucial role in bacterial physiology, survival, antimicrobial resistance and virulence (Heras *et al.*, 2009, Landeta *et al.*, 2018). The redox activities of archetypal Dsb proteins, best characterised in *E. coli* K-12, are typically tied to their oligomerisation state, i.e., monomeric and dimeric Dsb proteins, represented by DsbA and DsbC (and DsbG), which catalyse thiol oxidation and disulfide isomerisation, respectively (Heras *et al.*, 2009, Shouldice *et al.*, 2011, Landeta *et al.*, 2018). Much less is known, however, of other periplasmic thiol/disulfide oxidoreductases reported to adopt a variety of oligomeric structures and whether this can predict distinct catalytic activities in the cell remains unknown (Heras *et al.*, 2009).

Based on a growing body of evidence (Furlong *et al.*, 2017, Guddat *et al.*, 1998, Banaszak *et al.*, 2004, Bader *et al.*, 2001, Zhao *et al.*, 2003, Hiniker *et al.*, 2007, Santos-Martin *et al.*, 2021, Zapun *et al.*, 1995, Heras *et al.*, 2004), one hypothesis is that the functional fate of a thiol/disulfide oxidoreductase as an oxidase or reductase is likely to be governed by its oligomeric state. In this work, we directly investigated this question using a set of previously described ScsC proteins from different bacterial species, studied in the well-characterised genetic background of *E. coli* K-12. The three studied ScsC homologues were previously reported to adopt distinct quaternary architectures, including two recently characterised trimeric ScsC homologues (CcScsC and PmScsC) that are biochemically analogous to DsbC, and the monomeric StScsC that lacked disulfide oxidase and isomerase activity *in vitro* (Cho *et al.*, 2012, Petit *et al.*, 2022, Furlong *et al.*, 2017, Shepherd *et al.*, 2013, Furlong *et al.*, 2019). Here, we demonstrated that all three ScsC proteins exhibit dual dithiol oxidase and disulfide isomerase activities *in vivo*, irrespective of protein oligomerization status. Our findings may be explained by the natural limitations inherent to *in vitro* assays, in which the parameters may be affected by the artificial assay conditions that deviate from the cellular environment (Cho *et al.*, 2012, Petit *et al.*, 2022, Furlong *et al.*, 2017, Shepherd *et al.*, 2013, Furlong *et al.*, 2019) and highlight the importance of confirming enzyme function *in vivo*. The latter is of particular relevance to dithiol redox catalysts from diverse systems that are often found to co-exist in the same organism.

Bacterial periplasmic oxidoreductases are considerably diverse; while some are inherently housekeeping, others are only found in specific serovars or strains (Heras *et al.*, 2009, Landeta *et al.*, 2018). For instance, in addition to the housekeeping EcDsbA/EcDsbB pathway of *E. coli* K-12, uropathogenic *E. coli* strains also encode DsbL/DsbI, an extra redox pair that may be dedicated to the oxidative folding of virulence factors (Totsika, 2017). Cho *et al.* (2012) had reported the lack of a conventional Dsb reductive pathway in *C. crescentus* that is functionally supplied by CcScsC and CcScsB. However, in *S*. Typhimurium, the StScsC/StScsB pair co-exists with several Dsb oxidative and reductive pathways, and it was thought to exhibit no functional overlap (Shepherd *et al.*, 2013). Interestingly, we report that none of the characterised ScsC proteins can interact with EcDsbB, the cognate oxidase partner of DsbA in the prototypical *E. coli* system. However, consistent with previous biochemical evidence that StScsC and DsbD constitute a redox-relay pair (Subedi *et al.*, 2019), in our study, both CcScsC and StScsC, required DsbD to reduce disulfides efficiently *in vivo* (Fig. 3 and Fig. 4). Considering that DsbA/B and DsbC/D systems constitute the generic Dsb machinery, this fits the previous observation that reduced StScsC sequesters copper (Subedi *et al.*, 2019), i.e. reductive interactions with DsbD are, in fact, favourable for ScsC proteins, and explain the homology between ScsB (the cognate redox pair for ScsC) with DsbD, but oxidative reactions would deem detrimental to their capacity to bind copper (Cho *et al.*, 2012). Our finding further explained how ScsC proteins stay predominantly in their reduced form in the periplasm to counteract copper and oxidative stress (Subedi *et al.*, 2019, Shepherd *et al.*, 2013, López *et al.*,2018). Aside from pairing with ScsB/DsbD for redox relay, the ScsC proteins evade interactions with the prototypical Dsb oxidase system.

Interestingly, PmScsC still functions as an effective disulfide isomerase in the cell independently of DsbD, and a weak dithiol oxidase activity is observed despite the absence of a compatible re-oxidising partner in *E. coli* K-12. This observation could be explained by pools of reduced PmScsC being generated in the cell after having performed the oxidation function. It could also be proposed that yet unidentified partners are able to reduce PmScsC in the periplasmic space of *E. coli* K-12. Unlike the full-length PmScsC, the monomeric PmScsCΔN forms an oxidase pair with EcDsbB in *E. coli* and is functionally equivalent to EcDsbA (Fig. 2A), at the detriment of its original disulfide reductase activity (Fig. 5C). Thus, oligomerisation is adopted as a strategy to exclude undesirable oxidoreductive interactions. Bader *et al.* (2001) also showed that a monomeric variant of EcDsbC interacted with DsbB and acquired oxidase activity (Bader *et al.*, 2001).

However, upon trimerizing EcDsbA by engineering the N-terminal domain of PmScsC to the canonical EcDsbA, we observed no gain of disulfide isomerase function for this protein, and it retained oxidase activity *in vivo*. This could be due to the engineered EcDsbA units being in unfavourable orientations/positions within the chimera trimer to allow for productive interactions. Alternatively, the lack of isomerisation activity could be due to this activity not relying on trimerization of EcDsbA. Indeed, there is evidence of Dsb-like proteins not following the oligomerisation-isomerase rule, in that the monomeric StScsC was shown to be a suitable substrate of DsbD in the isomerisation pathway but not DsbB (Fig. 3 and Fig. 4). Overall, these findings suggest that there is more in guiding Dsb catalytic functions than oligomeric state alone. And while protein oligomerisation may serve as a mechanism to control functional overlap across different disulfide oxidoreductases in certain bacteria, we conclude that the cellular activities of periplasmic TRX-fold proteins cannot only be inferred by the protein’s oligomerisation status.

## AUTHOR CONTRIBUTION (CRediT)

YH: Conceptualisation, Methodology, Formal analysis, Investigation, Visualisation, Writing – Original and Draft, Writing – Review and Editing; JQ: Investigation, Writing – Review and Editing; LM: Investigation, Formal analysis, Visualisation; JJP: Investigation, Methodology, Writing – Review and Editing; BH: Visualisation, Supervision, Writing – Original and Draft, Writing – Review and Editing, Funding Acquisition; MT: Conceptualisation, Methodology, Supervision, Writing – Review and Editing; Funding Acquisition. All authors approved the final manuscript.

## ACKNOWLEDGEMENT

This work was supported by an Australian Research Council Discovery Project (DP190101613) to MT and BH; Clive and Vera Ramaciotti Health Investment Grant (2017HIG0119) to MT, and a Georgina Sweet Award for Women in Quantitative Biomedical Science to MT; The Ian Potter Foundation sponsored CLARIOStar high-performance microplate reader (BMG, Australia). MT received support from the Queensland University of Technology (QUT) through a Vice-Chancellor’s Research Fellowship. We thank Dr Anthony Verderosa for constructing pSU2718-EcDsbA. Vectors pMCSG7::PmscsC and pSC108 (DsbAss-CcScsC) were respectively gifted by Professor Jennifer L. Martin (Griffith University, Australia) and Professor Jean-François Collet (Institut de Duve, Universite’ Catholique de Louvain, Belgium). The authors thank Professor Jennifer L. Martin for critically reviewing the manuscript and acknowledge the use of the La Trobe University Comprehensive Proteomics Platform.

