## Supplemental Table S1 for "Bacterial suppressor-of-copper-sensitivity (Scs) proteins exhibit diverse thiol-disulfide oxidoreductase cellular functions"

| **Name** | **Sequence (5' to >3')** | **Detail** |
| --- | --- | --- |
| K-12 *dsbD* KO Forward | TTTTGATTTGTGTAAAGTTGAGATGCGTCTGAGTTTGCGCATGGGAATTAGCCATGGTCC | *dsbD* knockout forward primer |
| K-12 *dsbD* KO Reverse | ATTCGACGCGCCGGGACGTTCACAATTTGTCCCCGCGGATGTGTAGGCTGGAGCTGCTTC | *dsbD* knockout reverse primer |
| BamHI-*dsbC* Forward | TGCTCGGATCCATGAAGAAAGGTTTTATGT | *dsbC* cloning forward primer |
| HindIII-*dsbC* Reverse | ACTCGAAGCTTTTATTTACCGCTGGTCATTT | *dsbC* cloning reverse primer |
| KpnI-*dsbB* Forward | TTAGGGTACCATGATTATGTTGCGATTTTTGAACC | *dsbB* cloning forward primer |
| HindIII-*dsbB* Reverse | ATCTGAAGCTTTTAGCGACCGAACAGGTC | *dsbB* cloning reverse primer |
| BamHI-*dsbA* Forward | ATCAGGGTACCATGAAAAAGATTTGGCTGG | *dsbA* cloning forward primer |
| HindIII-*dsbA* Forward | TACGTAAGCTTTTATTTTTTCTCGGACAGATA | *dsbA* cloning reverse primer |
| HindIII-Cc*scsC* Reverse | CGATAAGCTTTCACCCCGCTTTGGCCCGCG | Cc*scsC* cloning reverse primer |
| PstI-*assT* Forward | ACGCTCTGCAGATGTTTGATAAATATAGAAA | *assT* cloning forward primer |
| KpnI-*assT* Reverse | CCAGGGTACCTTATTTGAACATCTGTTGTG | *assT* cloning reverse primer |
| Pm*scsC*^TD^ Forward | TGACATTGGAAGTGGATAAC | forward primer to amplify the trimerisation domain of Pm*scsC* |
| Pm*scsC*^TD^ Reverse | CAGTTTGGCATCTTTTGC | reverse primer to amplify the trimerisation domain of Pm*scsC* |
| *dsbA*^CAT^ Reverse | GCAAAAGATGCCAAACTGCAAGTGCTGGAGTTTTTCTCTTTC | forward primer to amplify the catalytic domain of *dsbA* |
| *dsbA*^CAT^ Reverse | TATCCACTTCCAATGTCATTTTTTCTCGGACAGATATTTCAC | reverse primer to amplify the catalytic domain of *dsbA* |

**Table S1. Oligonucleotides used in this study.**
